# Polymer brush bilayer at thermal equilibrium: A density functional theory approach

**DOI:** 10.1101/316141

**Authors:** Mike John Edwards

**Affiliations:** Leibniz Institute of Polymer Research Dresden

## Abstract

By means of the density functional theory (DFT) framework, the longstanding problem of the polymer brush bilayers at thermal equilibrium is studied. The calculated density profiles reveal that the brushes balance compression and interpenetration when they come into contact. The equation of state of the polymer brush bilayers is obtained and it represents scaling of the pressure with molecular parameters and distance between substrates. The results of this study may shed light in our understanding of some severe Musculoskeletal diseases which degrade the synovium. The significance of this study lays on the fact that the molecular structure is investigated through fundamental physical laws. So, this study bridges between theoretical biological physics and medicine.

## Introduction

Polymers are a class of long macromolecules that consist of repeated units of atoms or molecules i.e. monomers.^1,5,19^ Connectivity among monomers in a polymer chain is fulfilled through sharing valance electrons i.e. covalent or molecular bond. Monomers undergoing Brownian motions, constantly, change configuration of whole polymer chain so that in long time polymer chain undergoes all possible conformations. Polymer brushes are a class of polymeric structures that are present in biology, industry and medicine.^2^ Brush like structures could be found in outside of cell membranes. Polymer brushes form when linear chains are denesely grafted onto a surface. The chains strongly stretch in perpendicular direction due to the *steric repulsion* among monomers. Two opposing brush covered surfaces form a polymer brush bilayer (PBB). ^2^ PBBs are present in a wide range of biological systems such as mammalian knee joints as aggregan and capillaries of plants. ^2^ The most common knee joint disease which degrades severely the synovium (which is the thin layer of polymer brush that covers the joints.) is the synovial chondromatosis or synovial osteochondromatosis. In this disease, the swelling and inflammation of the synovium causes symptoms like joint stiffness. The exact underlying cause of synovium inflammation is unknown. Some researches suggest infection, trauma etc. The results of this research could shed light on causes of the synovial joint diseases through biophysical point of view. There have been numerous theoretical,^7–11^ experimental^15–17^ and numerical approaches to polymer brush bilayers. However, in the present article I approach the PBBs from an immensely different point of view. In the last decades, the theoretical approaches to PBBs have focused on the interpenetration zone between the opposing brushes. These theories are based on a concept which is called *blob picture*.^4,12–14^ The blob picture of the PBBs hypothetically says that the friction between two brush covered surfaces, originates solely from the interpenetration zone of brushes. In this article, I show that considering the structure of the whole system leads to correct equation of state for the PBBs based on the DFT framework.^6,18^ In the next section, the basics of the density functional theory (DFT) framework is introduced as a powerful theoretical tool in studying structure of biopolymers. In section, the DFT is applied to polymer brush bilayers and in the last sections, conclusions are discussed.

## Methods

The DFT framework is a useful theoretical tool in studying many body systems. Essentially, the DFT is based on describing free energy of physical systems in terms of their particle density and minimizing the grand potential functional with respect to density function and other parameters to find equilibrium properties. In systems with *inhomogeneous* density, the grand potential is a functional of density function. In such cases, *variational* derivative is applied to minimize the grand potential. Here, I describe constructing the grand potential of the polymer brushes in an step by step manner. As the first step, equilibrium properties of a polymer chain is discussed with help of *N-vector model*. Then, short-ranged pair interactions between monomers are discussed by using *Virial expansion* of the grand potential. At the end, the grand potential functional for polymer brushes is constructed and discussed.

### Polymer chain in an external force

Conformation of a polymer chain constantly changes due to the *Brownian motion* of its monomers. Theoretically, this condition could be assumed as *N* vectors of length *a* which are connected to each other and are freely rotating in three dimensional space. To obtain elastic properties of the chain, we need to apply an external force on it and observe its response. At the end, one has to remove the external force from the resulting expressions. The remaining terms represent elastic properties of chain in absence of the external force. First of all, I introduce the *Hamiltonian* of the system as a summation over inner products of the external force with all vectors as follows,

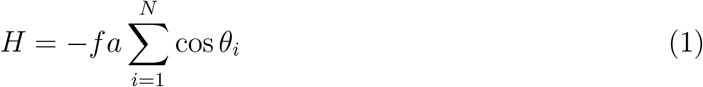

Negative sign here ensures that minimum energy takes place when all vectors are directed to the direction of external force. So, conformations of a polymer chain are characterized by *θ*, the smaller angle between the external force and vectors. For simplicity and symmetry considerations, the spherical polar coordinates are chosen and the external force is applied in the z direction. This way, the angle between the external force and each vector is the same as the polar angle *θ* in the spherical polar coordinates. Having introduced the Hamiltonian, the next step would be calculation of the partition function which is defined as *Z* = ∫ *d*Ωexp – *H*/*k*_B_*T* where *d*Ω = *d**x***_1_ … *d**x**_N_* denotes a hyper volume element in configuration space of *N* vectors. The partition function reduces to the following integral in the spherical polar coordinates,

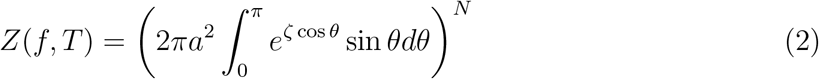

where *ζ* = *af*/*k*_B_*T* is a dimensionless quantity. The above integral is simply solvable due to the fact that the vectors (monomers) are uncorrelated and it converges to the following result for the partition function,

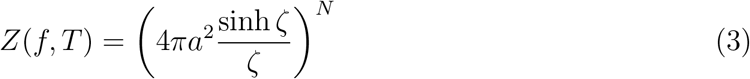

Now we can simply calculate the canonical free energy *F*(*f, T*) = –*k*_B_*T* log *Z*(*f, T*) as follows,

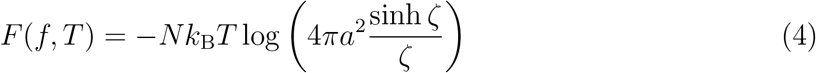

This is free energy in external force and temperature ensemble. Since, the external force and temperature are fixed, so the chain length fluctuates. Expectation value of the chain length can be obtained from canonical free energy by –*∂F*(*f, T*)/*∂f*,

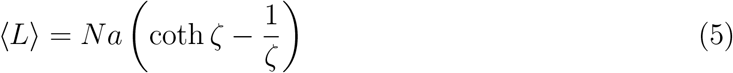

Expectation value of the chain length could alternatively be calculated from direct integration which gives us the above result as well. Note that, in statistical mechanics, coth *ζ* – *ζ*^−1^ is called the *Langevin function* and it extensively is used.^3^ It turns out that, expectation value of the chain length subject to an external force is non-zero. Certainly, this occurs, exclusively, along the external force direction (here the z direction). In other directions (here x and y directions) expectation value of the chain length vanishes. To obtain fluctuations of the chain length, one needs to have expectation value of the mean-squared-length i.e. 〈*L*^2^〉. Expectation value of the mean-squared-length could be calculated through direct integration leading to the following result,

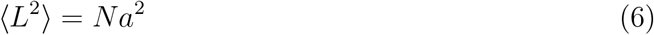

and fluctuation in the chain length is calculated via 〈*L*^2^〉 – 〈*L*〉^2^ as,

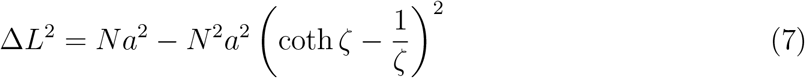

Susceptibility *∂*^2^*F*(*f, T*)/*∂f*^2^ which is a measure of fluctuations reads as follows,

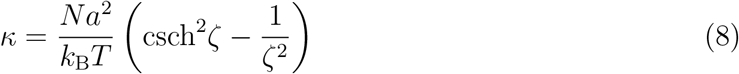

In the DFT framework, response of the chain in the length and temperature ensemble is essentially considered. This could be obtained by calculating the Helmholtz free energy via *Legendre transformation A*(*L, T*) = *F*(*f, T*) + *f*〈*L*〉. The Helmholtz free energy of a polymer chain is given as follows,

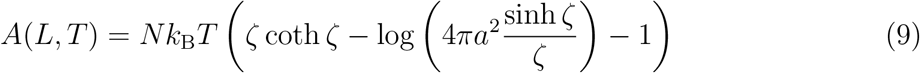

### Polymer chain in an strong external force

A interesting limiting case, is a very strong external force applied to a polymer chain. In this extreme limit, the chain stretches strongly in direction of the external force and the chain reaches the rod limit at sufficiently large external forces. The behavior of physical quantities under this extreme condition could be looked at by making limit of them when the external force approaches infinity as follows,

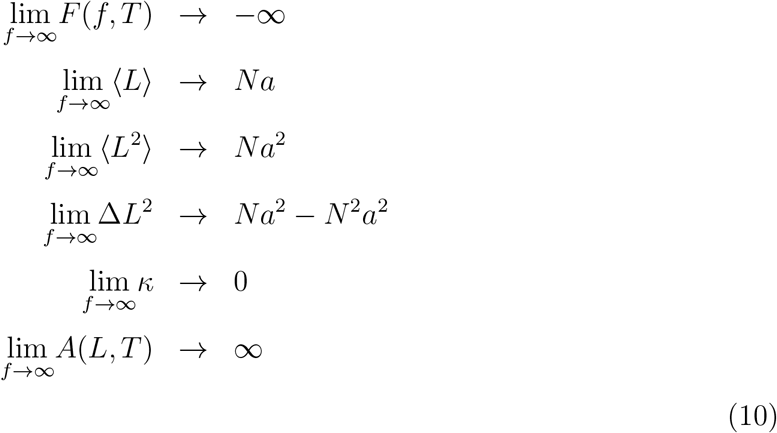

In the rod limit, the expectation value of the chain length goes to *Na*. The squared fluctuations goes to *Na*^2^ – *N*^2^*a*^2^ as the rod limit is achieved, however, this is a negative value hence leads to an imaginary quantity for the fluctuations. Therefore, one could argue that at the rod limit, fluctuations is not defined. The susceptibility goes to zero i.e. the elasticity vanishes which means that the chain becomes a very stiff spring. The free energies go to infinity at the rod limit stating that there is no fluctuations.

### Polymer chain in a weak external force

In most biological systems, magnitude of the external force is much smaller than the thermal energy. In this case, one could consider the dimensionless quantity *ζ* ≪ 1 and make the Taylor expansion over all physical quantities. The Taylor expansions give us the following terms,

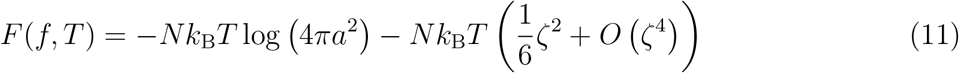

The first term in the canonical free energy is free energy of *N* freely rotating vectors in absence of the external force. The second term is the leading order term of the canonical free energy which contains the external force. This term has the most of information about elasticity of a polymer chain.

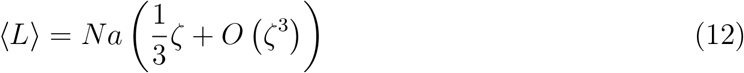

In Eqs. 12 and 13 average length and *compressiblity* of the chain is seen in a weak external force. The leading order term in average length depends on the external force so in absence of the external force, the average length vanishes, as it is expected. Naturally, when a polymer chain is subject to an external force, it tends to direct itself to the external force. This causes non-zero average length direction of the external force.

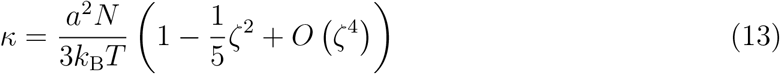

The leading order term of compressibility is independent of external force. It gives an intrinsic property of polymer chain 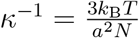 which indicates the elasticity originated from entropy. By looking at Eqs. 12 and 13, we see that the leading order term in the canonical free energy could be rewritten as 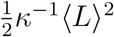. This free energy shows the *entropic elasticity* in a polymer chain. The entropic elasticity tries to shrink the chain. However, it is different from crystals. The elastic constant in crystalline solids decreases by temperature, however, in a polymer chain and rubber-like materials the elastic constant increases by temperature. So, by heating up the polymers they contract due to 〈*L*〈 = *f/κ*^-1^ term in contrast to the crystalline materials. One could easily check this physical law by heating up a nylon bag.

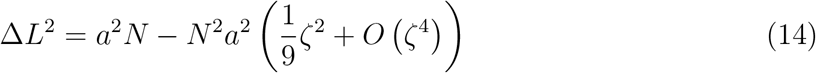

Fluctuation of chain length in weak external force is extremely interesting quantity. To find the leading order term in the perturbation expansion, we need to compare their orders of magnitude. Since the second term scales as ~ *N*^2^, it appears to dominate over the first term which scales as ~ *N*. Nevertheless, the second term is multiplied by *ζ*^2^. So to find the leading order term, one needs to make numerical comparison. Typical polymer chains consist of *N* = 10^5^ monomers and usually force is in order of pN, and typically, monomer size is 0.1 nm. Considering order of magnitudes of the Boltzmann constant as 10^−23^ and room temperature *T* = 300K, the perturbation coefficient becomes *ζ* ≈ 10^−2^. Consequently, the first and the second terms have almost the same orders of magnitude and both of them are considered as leading order terms. As it is already expected, when temperature increases, fluctuations of the chain length increases as well and in the extreme limit of *T* → ∞, we have Δ*L*^2^ → *a*^2^*N*. It means than fluctuations of the chain length equals to the expectation value of the squared chain length. In presence of the external force, fluctuations are small, as compared to the expectation value of the squared chain length. In absence of the external force, fluctuations equal to 〈*L*^2^〉. It means that under influence of the external force, the chain is stretched in the external force direction and shows minor fluctuations in length. The Helmholtz free energy at small external forces is given as follows,

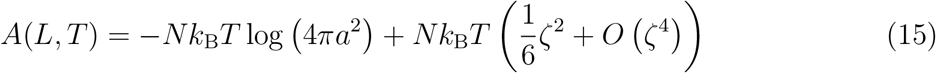

It shows that elasticity of a polymer chain is *κ*^−1^〈*L*〉^2^/2 at small external forces. This is an significant result which will be used in the DFT framework as the grand potential of a polymer chain.

### Short-ranged interactions between monomers

Now assume *N* particles that can freely move and interact with each other through short-ranged forces. The Hamiltonian of such a system is given as *H* = *K* + *U* with *K* total kinetic energy and *U* total potential energy of the system. The most simple case is the binary interactions. In such a particular case, total potential energy becomes a summation over all pair potentials. The total potential energy is a summation over *N*(*N* – 1)/2 possible pair potentials which excludes repeated and self interactions. In systems that are exposed to no external force and have *translational invariance*, the pair potential is only a function radial distance between particles. In such an special case, one can write down the canonical partition function of the system as follows,

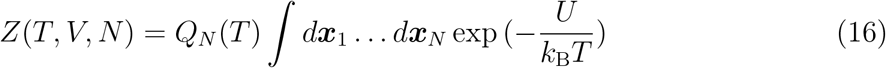

Calculations show that the above partition function would lead to the *Virial expansion* of the Helmholtz free energy which one would write it down as follows, ^3^

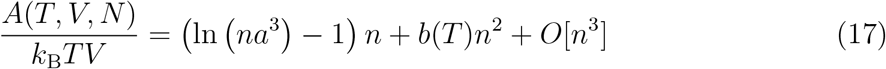

Coefficient of *n*^2^ in Eq. (17) is called the *second Virial coefficient* which by definition is given as follows,

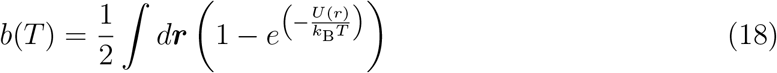

The second Virial coefficient is key quantity in describing short-ranged binary interactions between particles. For hard spheres of diameter *a*, *b* = (2*π*/3)*a*^3^ and is independent of temperature. For soft spheres, the magnitude of the second Virial coefficient gets something larger than hard spheres. However, more realistic pair interaction which has been observed at molecular and atomic scales is the Lennard-Jones pair potential which reads as *U*(*r*) = 4*ϵ*[(*a/r*)^12^ – (*a/r*)^6^]. The Lennard-Jones pair potential is extensively used in coarse-grained computer simulations. For a Lennard-Jones pair potential, one could precisely calculate the second Virial coefficient at low temperatures limit as follows,^3^

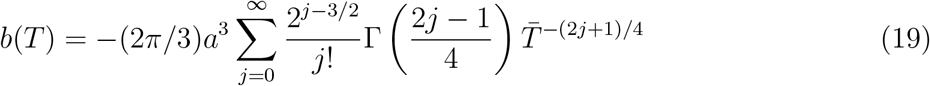

with 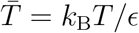 defined as a dimensionless quantity.^3^ For instance, if 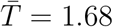, then the second Virial coefficient converges to *b* = −2.

In polymer solutions, a positive b refers to *good solvent* conditions and a negative b refers to *poor solvent* conditions. This definition is based on the fact that, monomers are treated much larger than solvent molecules and information about solvent molecules is contained in b, effectively. However, when solvent molecules are comparable with monomers, one has to take the interactions between solvent and monomers explicitly. Additionally, at a certain temperature (*θ* temperature) the second Virial coefficient vanishes. At *θ* temperature, monomers do not make pair-wise interactions, however, the higher order interactions take place.

### Polymer brushes

Polymer brushes could be tackled by the DFT framework.^6,18^ Consider linear polymer chains that are grafted to a substrate at *z* = 0 (See Fig. (1)). Number of grafted chains per surface area is defined as the grafting density *σ*. Since, one deals with an in-homogeneous system, the grand potential possesses a functional form of the density function. The grand potential functional of the polymer brushes is given as follows,

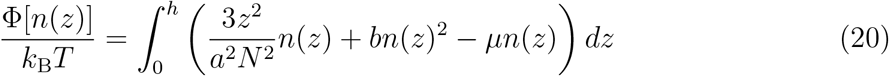

**Figure 1:**
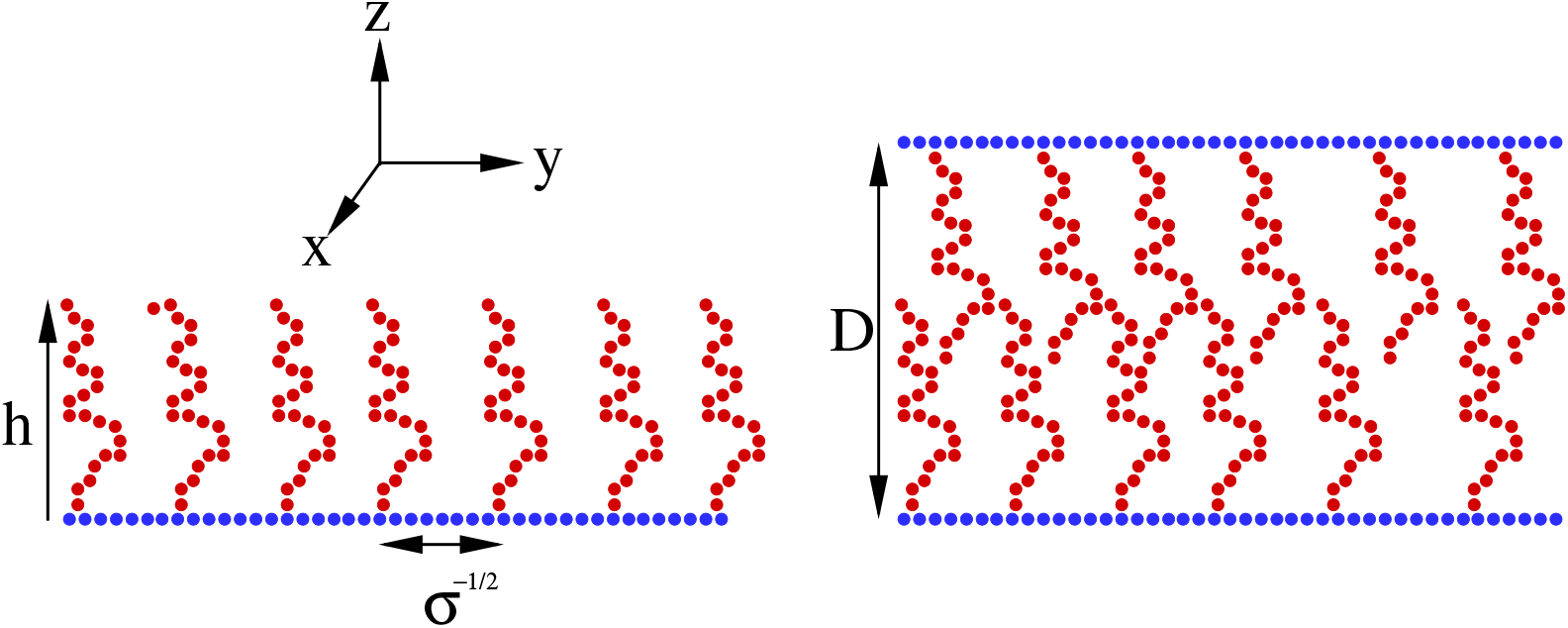
Schematic view of a polymer brush (left) and a polymer brush bilayer (right).

The first term in Eq. 20 refers to leading order term of the Helmholtz free energy which describes the entropic elasticity of chains. Since energy is distributed throughout the chain according to the probability distribution function, one could multiply the Helmholtz free energy by *P*(*z*) = *n*(*z*)/*N* with *n*(*z*) the density profile of monomers. The second term refers to the Virial expansion of the grand potential functional and it takes the excluded volume interactions among monomers into account. The third term applies a constraint over entire system to keep total number of monomers fixed with *μ* denoting the chemical potential or the *Lagrange multiplier*.

The equilibrium density profile, brush height and chemical potential are obtained through simultaneously solving the following set of coupled equations. The first equation sets a functional derivative of grand potential with respect to the monomer density profile. It leads to an Euler-Lagrange equation. Solution of the corresponding Euler-Lagrange equation gives us the equilibrium density profile. The second equation refers to derivative of grand potential with respect to brush height including the obtained equilibrium density profile. The third equation indicates that total number of monomers in brush is obtained through integration over density profile throughout brush region.

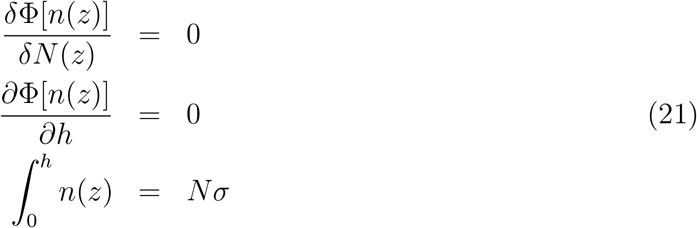

Simultaneously solving the set of coupled equations at Eqs. 21, leads to the following results,

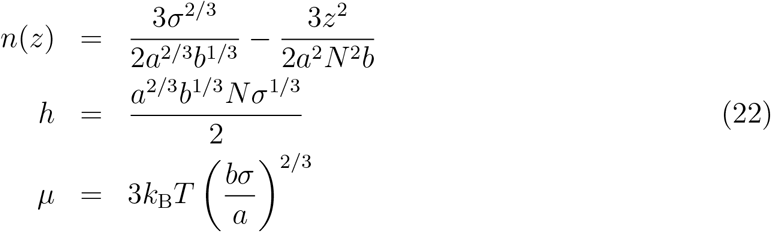

Eq. 22 reveals a parabolic density profile for brush as well as universal power laws for brush height and chemical potential in terms of molecular parameters. Inserting Eq. (26) into Eq. (21) gives us grand potential in terms of molecular parameters as follows,

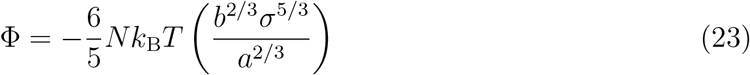

which indicates that grand potential scales with molecular parameters as well.

## Results

### Polymer brush bilayers

The most designated application for the PBBs is devoted to lubrication inside mammalian Synovial joints. In such a case, two opposing brush covered substrates are at a distance where brushes intermediately compressed over each other. Here, I consider two opposing brush covered surfaces at *z* = 0 and *z* = *D* (See Fig. (1)). The following grand potential seems to be appropriate for this problem where the first integral represents the bottom brush, the second integral represents the top brush and the third integral represents the steric repulsion among monomers of two brushes throughout interpenetration zone. The two brushes are exactly the same as each other in molecular parameters, so, to take the second brush into account, it just suffices to transform *z* → (*z* – *D*) in entropic elasticity term plus integrating from height of the top brush (*D* – *h*) to *D*. To take steric repulsion among two brush layers into account, one could integrate over a multiplication of both density profiles within the interpenetration zone i.e. from (*D* – *h*) to *h*.

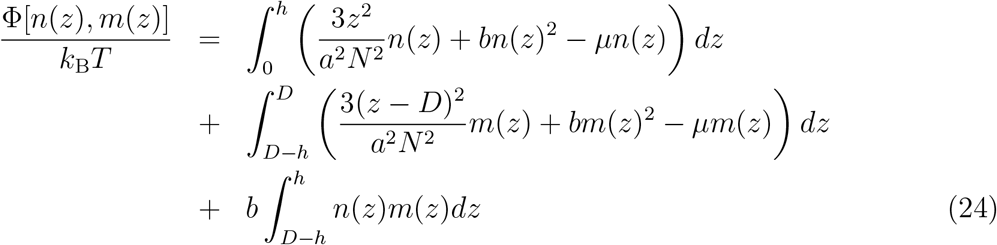

Having introduced the grand potential functional, solving the following equations lead to equilibrium density profiles, brush height and chemical potential corresponding to each brush.

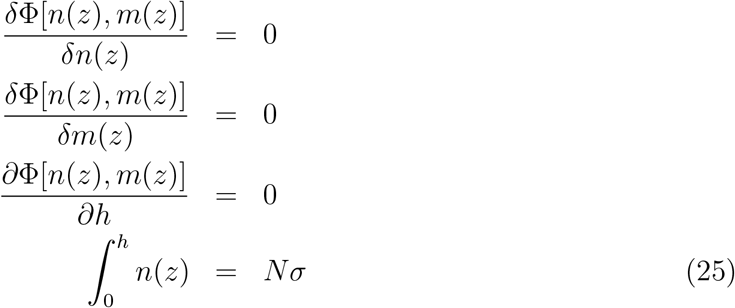

Note here that, it suffices to take derivative of grand potential functional only with respect to *h* as well as to calculate total monomers of only bottom brush in calculations. This becomes possible due to similarity of two brushes. Eqs. 25 lead to the following results for monomer density profiles, brush heights and chemical potential. Note that, height of the top brush is calculated as (*D – h*).

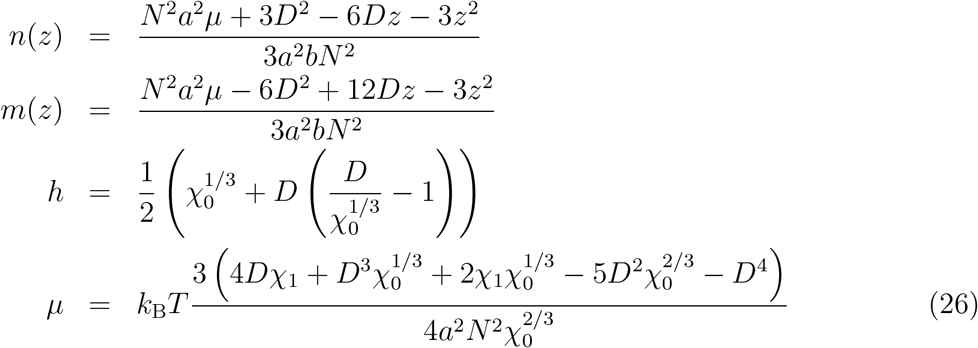

where χ_0_ and χ_1_ are volume scales as defined here,

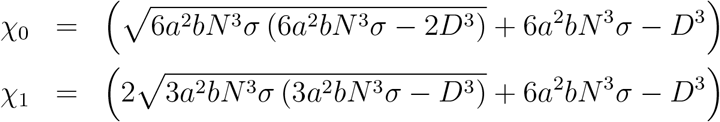

Now we can calculate the interpenetration length between two brushes through calculating 2*h* – *D*. This leads to the following result,

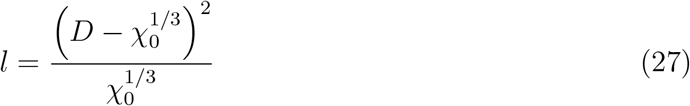

In Fig. 2, the density profiles of brushes, their overlap density profile are presented. The molecular parameters are chosen as *a* = 1, *b* = 2.09, *σ* = 0.1, *N* = 30 and *D* = 20. Density profiles of brushes at intermediate compression possesses three major properties; larger density of monomers in vicinity of surfaces, smaller thickness and non-parabolic (almost linear) profile in perpendicular direction. These are considered as response of brushes to an intermediate compression through balancing *compressive* forces by interpenetration.

**Figure 2:**
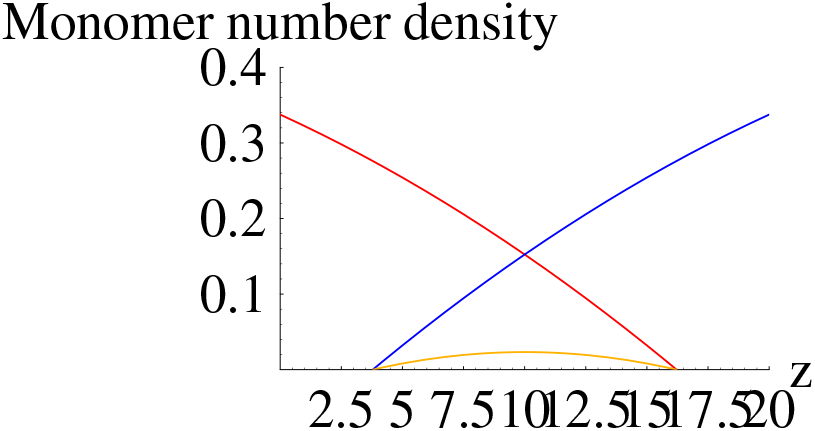
Density profiles of left brush (blue), right brush (orange), the overlap density profile of left and right brushes (green) and density profile of left brush in absence of right brush (red).

The brush height in terms of molecular parameters is shown in Fig. (3). It turns out that, the brush height scales as *Nσ*^0.38^*b*^0.4^. At small segment lengths and wall distances, the brush height scales as *a*^0.88^*D*^−0.1^, however, at large segment lengths and wall distances, it scales as *a*^0.69^*D*^−0.5^.

**Figure 3:**
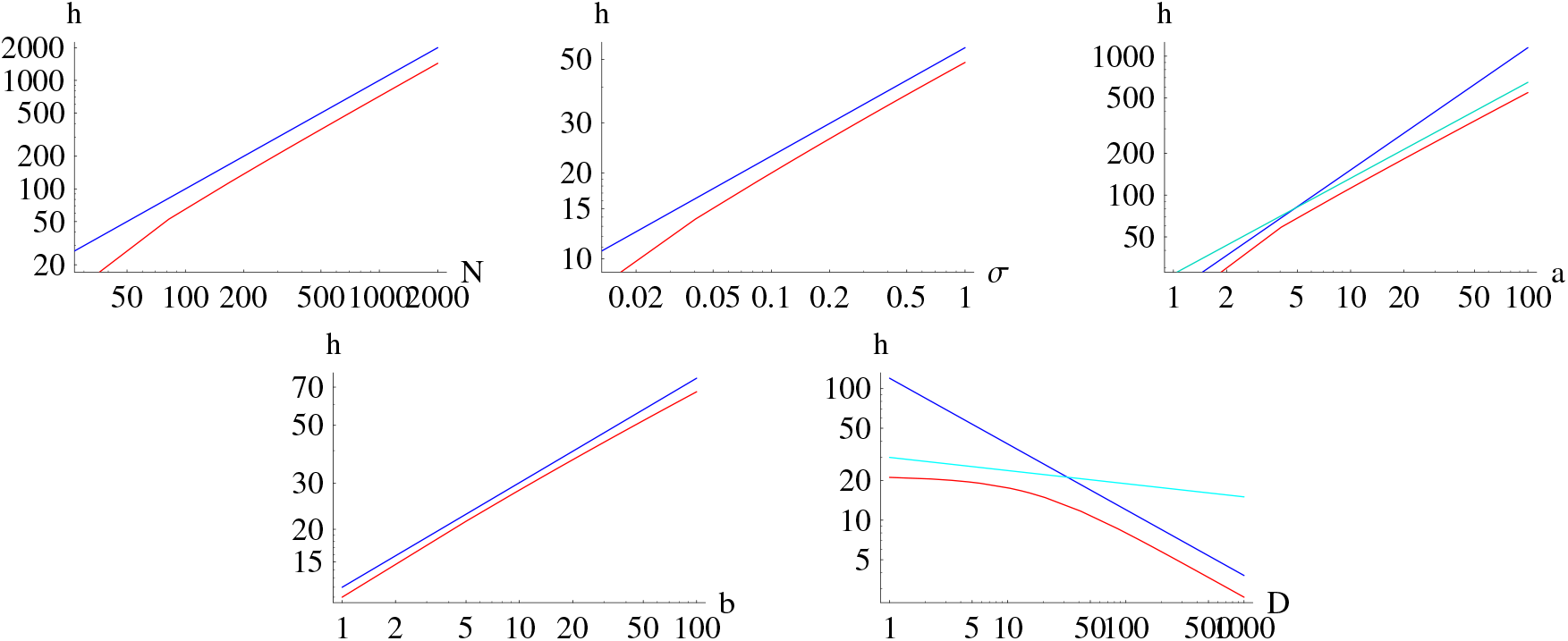
The brush height in terms of the molecular parameters of the system. The red lines show the DFT result and blue lines show the fitted power laws. It turns out that the brush height scales as *Nσ*^0.38^*b*^0.4^. At low segment size and wall distance, the brush height scales as *a*^0.88^*D*^−0.1^ and at high segment length and wall distance, it scales as *a*^0.69^*D*^−0.5^.

In Fig. 4, interpenetration length in terms of wall distance is presented. This plot, in contrast to what has been claimed in works already published, does not fit to any universal power law. However, interpenetration length decreases with increasing wall distance which is already expected. In Ref., ^4^ it is mentioned that the interpenetration length in polymer brush bilayers at melt concentration scales as ~ *D*^−1/3^ and at semidilute concentration as ~ *D*^−0.17^. However, the wall distance appears not to follow any power law. The interpenetration length scales as ~ *N*. This power law almost fits to Eq. 27. In Ref.,^4^ it is argued that the interpenetration length scales as ~ *N*^2/3^ and ~ *N*^0.51^ respectively for melt and semidilute regimes. Naturally, as degree of polymerization increases, longer polymer chains tend to increase interpenetration length. Furthermore, the chains inside the interpenetration zone are not folded back. To this reason, the interpenetration length scales as ~ *N*. The second Virial coefficient scales as ~ *b*^0.35^. It is natural that bilayer respond to binary correlations among monomers. For instance, quality of solvent is determined by the second Virial coefficient as poor, theta or good solvents are considered. The interpenetration length scales as ~ *σ*^0.34^. In Ref., ^4^ the interpenetration length is independent of the grafting density for melts and it scales as ~ *σ*^−0.51^ for semidilutes. Nevertheless, one would criticize the results of Ref.^4^ since the interpenetration length naturally must increase with number of chains per surface area. The interpenetration length scales as ~ *a*^0.67^ while Ref.^4^ proposes a power law ~ *a*^4/3^ for melt and ~ *a*^0.15^ for semidilute PBBs.

**Figure 4:**
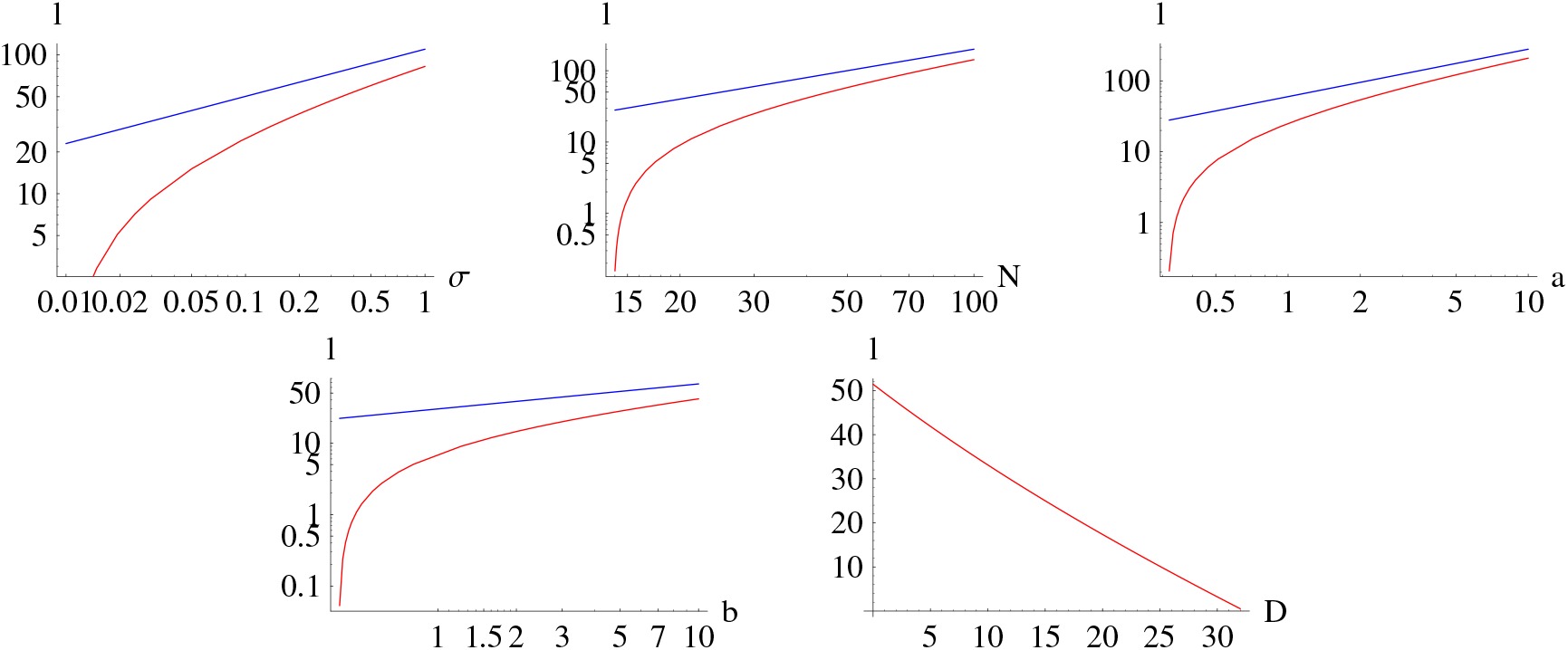
Interpenetration length in terms of the molecular parameters of the system. The red lines show the DFT result and blue lines show the fitted power laws. The plots show that the interpenetration length scales as *σ*^0.34^*a*^0.67^*b*^0.35^*N*. However, there is no power law in terms of the wall distance.

The total grand potential can be calculated by inserting Eq. (26) into the grand potential. By integration we obtain,

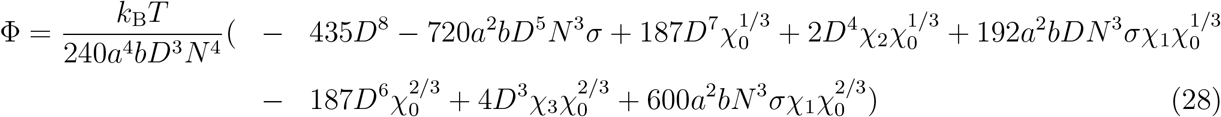

where the volumes are defined as follows,

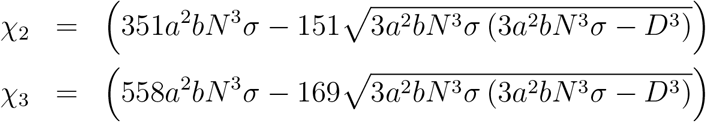

In Fig. (5), Eq. (28) is plotted in terms of molecular parameters of the system. It turns out that, the grand potential scales as *σ*^2.7^*b*^1.7^*a*^1.3^*N*^4^*D*^−3^. In Fig. (6), the grand potential landscapes has been plotted in terms of molecular parameters of the system. They are plotted by choosing molecular parameters as *a* =1, *b* = 2.09, *σ* = 0.1, *N* = 30 and *D* = 20. The grand potential landscapes are useful in checking the stable and metastable points in the pressure. To calculate pressure, one could use the Gibbs-Duhem relation *P* = –Φ/*V*. Since, the system is at thermal equilibrium, the pressure is the same as throughout the medium. The pressure as well as the equation of state is given as follows,

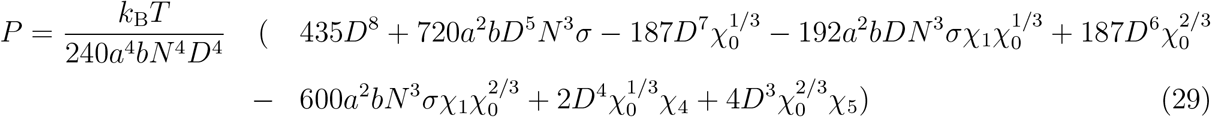

where the volumes χ_4_ and χ_5_ are defined as follows,

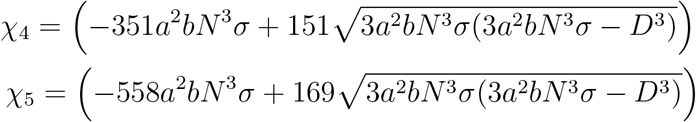

**Figure 5:**
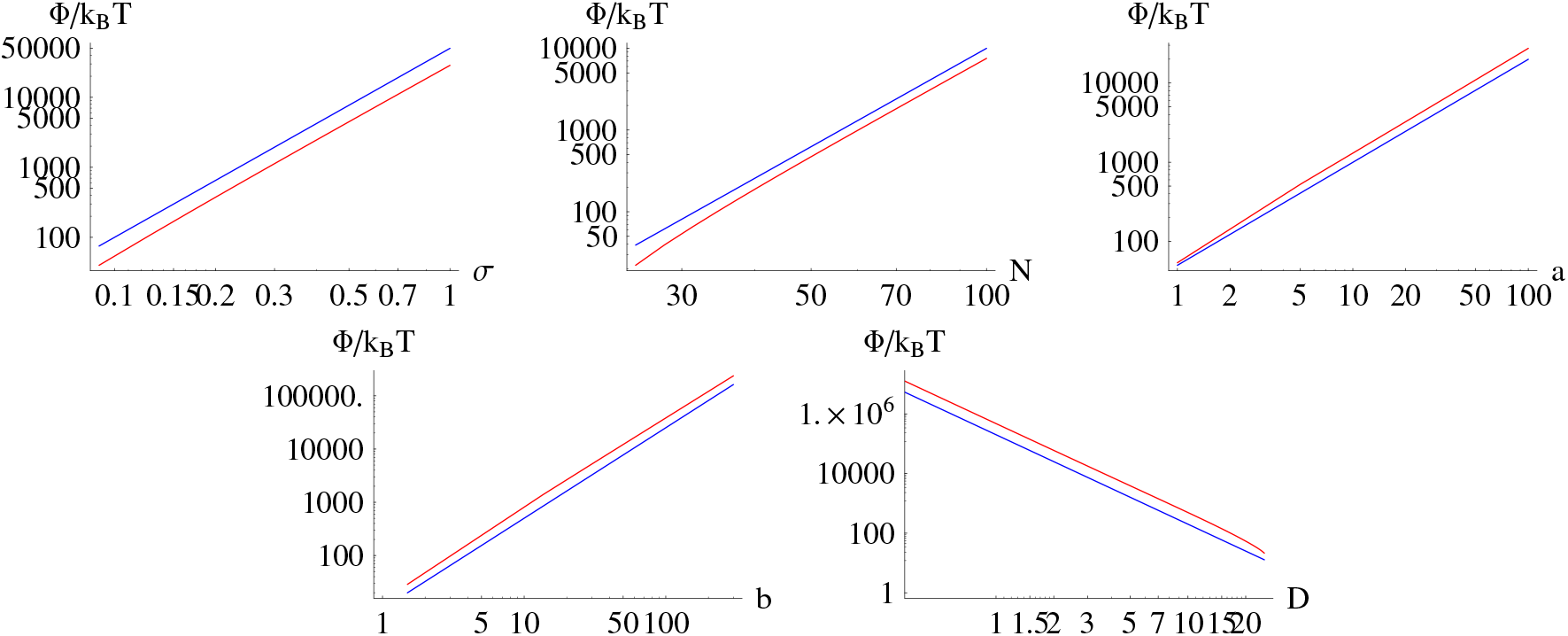
Total grand potential in terms of the molecular parameters of the system. The red lines show the DFT result and the blue lines show the fitted power laws. It turns out that the grand potential scales as *σ*^2.7^*b*^1.7^*a*^1.3^*N*^4^*D*^−3^

**Figure 6:**
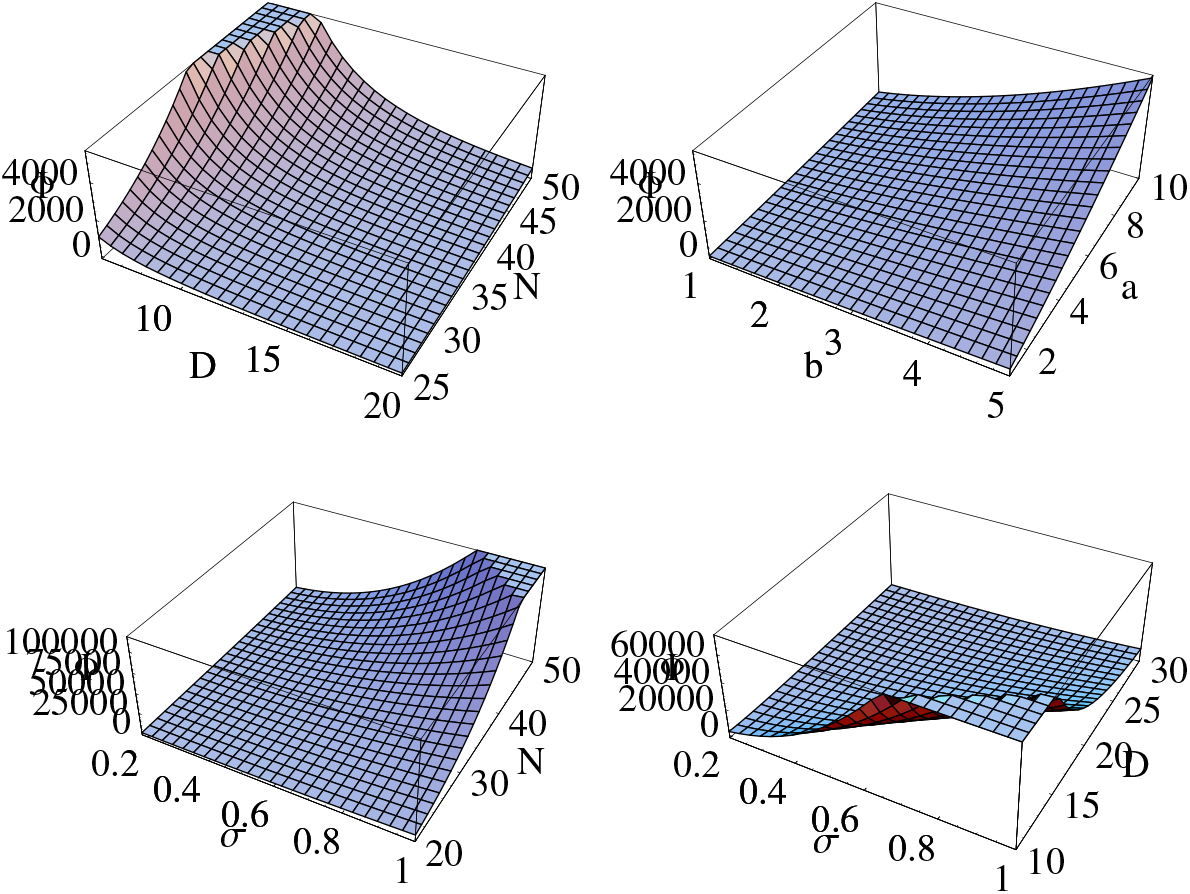
The grand potential landscapes in terms of molecular parameters of the system.

In Fig. (7), Eq. (29) in terms of the molecular parameters of the system is plotted. It turns out that the pressure scales as,

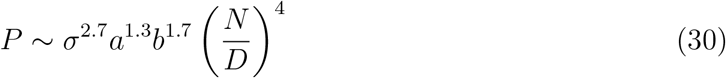

**Figure 7:**
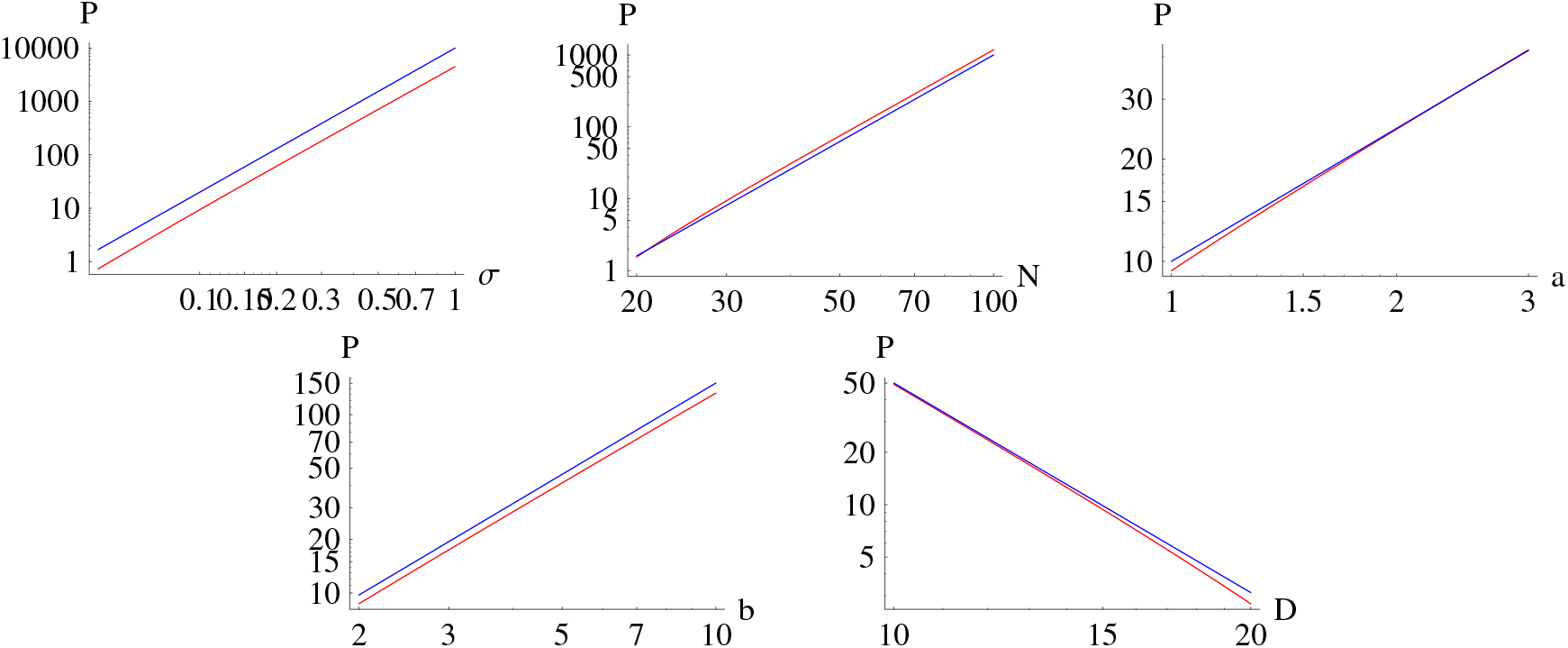
The pressure exerted on the bottom substrate in terms of the molecular parameters of the system. The red lines show the DFT result and the blue lines show the fitted power laws. It turns out that, the pressure scales as *σ*^2.7^*a*^1.3^*b*^1.7^*N*^4^*D*^−4^.

## Discussion

The results of the DFT framework shows that the density profile of brushes in a PBB do not follow the parabolic profile of an isolated brush. Instead, curvature of the PBB density profile is much less than an isolated brush. This reveals influence of compression and interpenetration on the distribution of monomers. This is important because it discloses information about distribution of monomers throughout a PBB. The brush height scales as *N* even in the PBB. This tells us that the chains reside in the strong stretching limit (SSL) despite compressive forces from the opposing brush. Additionally, the brush height slightly decreases by the wall distance D. This result emphasizes that the chains tend to shrink at larger wall distances *D*. This is opposite to what might be thought at first view. The interpenetration length decreases by the wall distance. This result tells us that the brushes start reduce interpenetration when wall distance increases. The DFT framework tells us that all molecular parameters contribute to increase the interpenetration length. The grand potential shows a perfect power law behavior. The grand potential landscapes show a very smooth landscape in terms of *N* and *D*, *N* and *σ* as well as *a* and *b*. The pressure decreases by the wall distance as *D*^−4^ which is natural result of reduction in strong compressive forces. This power law is a correction to the previously reported *D*^−3.5^ in the work already published^4^ and others. The equation of state of the PBBs obtained here proofs a very important point and that is the fact that taking into account only binary interactions leads to the correct equation of state even in very compressed situations. This fabulous fact proofs that binary interactions play the most important role in the monomer interactions.

## Conclusion

In this study, the DFT framework is made to approach the polymer brush bilayers at equilibrium conditions. The framework shows how polymer brushes interpenetrate into each other through giving density profiles of two brushes. It also shows how the interpenetration length depends upon molecular parameters and wall distance. The interpenetration length decreases by wall distance through a non-power law behavior. In contrast to previous publications, this suggests that the interpenetration length does not obey similarity rules in terms of wall distance. Although, it scales with molecular parameters. Significance of this study becomes more clear as it is a unique approach to problem of interpenetration of two soft materials by taking their simultaneous deformations into account. Simultaneous deformation is a key element in approaching soft matter and biological systems. The blob picture which have been utilized in Ref.^4^ seems to be extremely simple and it does not capture important details. For instance, interpenetration length is dependent on binary correlations among monomers as well as it increases by grafting density, two facts that blob picture does not take into account. It is worth noting that since in biological macromolecules there are competitions among different interactions (such as elasticity, excluded volume etc), therefore approaches like blob picture are not able, intrinsically, to deliver sufficient information. Regarding the application of these results in synovium diseases, it is important to know that the inflammation and swelling of synovium tissue occurs due to many parameters. For instance, the DFT framework reveals that the chains swell by all molecular parameters. The wall distance shrinks the chains. So, one could argue that reduction in the bones distance inside the knee joints, would lead to swelling.

## Acknowledgments

This is my great honor to thank the Cold Spring Harbor Laboratory (CSHL) and the Regeneron Pharmaceuticals Inc. for financial support, Wolfram Research for Mathematica, the Red hat Inc. for Fedora Linux, Tecplot Inc. for Tecplot visualization software, Weizmann Institute of Science for XMGrace.

